# Shifts in attention drive context-dependent subspace encoding in anterior cingulate cortex during decision making

**DOI:** 10.1101/2023.10.10.561737

**Authors:** Márton Albert Hajnal, Duy Tran, Zsombor Szabó, Andrea Albert, Karen Safaryan, Michael Einstein, Mauricio Vallejo Martelo, Pierre-Olivier Polack, Peyman Golshani, Gergő Orbán

## Abstract

Attention is a cognitive faculty that selects part of a larger set of percepts, driven by cues such as stimulus saliency, internal goals or priors. The enhancement of the attended representation and inhibition of distractors have been proposed as potential neural mechanisms driving this selection process. Yet, how attention operates when the cue has to be internally constructed from conflicting stimuli, decision rules, and reward contingencies, is less understood. Here we recorded from populations of neurons in the anterior cingulate cortex (ACC), an area implicated in ongoing error monitoring and correction during decision conflicts, in a challenging attention-shifting task. In this task, mice had to attend to the rewarded modality when presented identical auditory and visual stimuli in two contexts without direct external cues. In the ACC, the irrelevant stimulus continuously became less decodable than the relevant stimulus as the trial progressed to the decision point. This contrasted strongly with our previous findings in V1 where both relevant and irrelevant stimuli were equally decodable throughout the trial. Using analytical tools and a recurrent neural network (RNN) model, we found that the linearly independent representation of stimulus modalities in ACC was well suited to context-gated suppression of a stimulus modality. We demonstrated that the feedback structure of lateral connections in the RNN consisted of excitatory interactions between cell ensembles representing the same modality and mutual inhibition between cell ensembles representing distinct stimulus modalities. Using this RNN model showing signatures of context-gated suppression, we predicted that the level of contextual modulation of individual neurons should be correlated with their relative responsiveness to the two stimulus modalities used in the task. We verified this prediction in recordings from ACC neurons but not from recordings from V1 neurons. Therefore, ACC effectively operates on low-dimensional neuronal subspaces to combine stimulus related information with internal cues to drive actions under conflict.

## Introduction

With a limited capacity to process sensory stimuli in an information-rich world, how the brain distinguishes relevant from irrelevant stimuli is one of the core problems in neuroscience. Out of a torrent of sensory features the nervous system is required to select the limited few that predict the outcome of actions. By learning to associate specific cues to specific task contingencies, animals can adapt to situations in which the relevance of features can change. In such cases, learning reduces to a simple conditioning paradigm, where cues determine the set of features attended. One proposed mechanism of attention is the enhancement of the relevant and suppression of the irrelevant feature (Kaping et al., 2011). However, a fundamentally different computational problem arises when there are no cues to unambiguously establish the set of features attended or when relevance of cues is volatile (Heald et al., 2021). In these conditions, it is unclear how conflicting contingencies between stimulus and outcome can be resolved by attention.

To address this question, we adopted a set-shifting task (Birrell & Brown, 2000; Hajnal et al., 2021; Spellman et al., 2021) in which animals were presented with both visual and auditory stimuli, of which only one was relevant for obtaining a water reward, while the other acted as a distractor. Whether visual or auditory stimuli were relevant for obtaining reward was consistent during a block of trials and was switched at block boundaries. In this set-shifting task no external cues were provided to signal the actual relevance of the presented stimuli. Consequently, the task required the animals to cope with the changing relevance of stimulus modalities. Thus, in the absence of direct cues, animals inferred the contexts in order to resolve apparent conflicts by constructing an internal representation from stimulus, decision, and reward contingencies. How the cortical circuits can support this computation has remained unclear.

The anterior cingulate cortex (ACC) has long been implicated in monitoring conflicts (Botvinick et al., 1999). More recently, it has been proposed that ACC plays a more active role in updating stimulus-outcome contingencies, modulating its activity after attentional errors (Shen et al., 2015), modifying behavior after negative experiences (Oemisch et al., 2019) and improving value estimates for planning goal oriented behaviour (Hayden et al., 2011; Holroyd & Yeung, 2012; Kennerley et al., 2006; Silvetti et al., 2013). Therefore, during the set-shifting task, while prefrontal cortex (PFC) collects action and outcome history (Seo and Lee, 2017), ACC is modulated by past information when conflicts induce errors (Dosenbach et al., 2006; Kennerley et al., 2006; Seo & Lee, 2007). Without a cue indicating changes in stimulus-outcome contingencies, i.e. in the relevance of particular stimuli, even after learning the paradigm, the animal may rely on active conflict monitoring to establish an internal cue that identifies the currently relevant features. Notably, active conflict monitoring is critical before completely learning the behavioral paradigm and therefore ACC is expected to have an important role during learning as well. Taken together, we hypothesize that by tracking conflicting situations, ACC generates internal cues that define the relevance of sensory features. Then, relying on these cues, ACC implements a context-gated attention mechanism that prioritizes information relevant for the outcome of actions by suppressing the irrelevant input.

To understand how attention mechanisms select the relevant features in a given context, we examined the low dimensional representations of neural activity in ACC during performance of the set-shifting task. We found that neural correlates of stimulus and context are represented in geometrically interpretable low-dimensional manifolds, often in linear subspaces, as previously observed (Bernardi et al., 2020; Hajnal et al., 2021). We found that in this subspace the irrelevant stimulus modality was systematically suppressed during sensory stimulus presentation. We identified context-dependent gating of the irrelevant stimulus as a mechanism that is well aligned with the representation found in ACC. Using a Recurrent Neural Network (RNN) model of the set-shifting task, we reproduced key properties of our ACC recordings. A signature of context-dependent gating was demonstrated in the RNN, which was then later identified in neural recordings as well, attesting to the utility of RNNs informing analysis of experimental data. Our work thus provides a mechanistic account of how stimulus relevance can determine our choices under conflicting evidence.

## Results

To dissect the neural mechanisms of attentional set-shifting we trained mice to perform an audio-visual cross-modal context-dependent decision making task (Hajnal et al., 2021). In this task we simultaneously and randomly displayed one of two visual stimuli (45 degree or 135 degree drifting gratings) and one of two auditory stimuli (5000 and 10000 Hz tones). During each block of trials mice had to make Go/No-go decisions (licking for water reward) based on one sensory modality while ignoring the other modality. Therefore, only one of the stimulus modalities was relevant for making a correct decision, while the other modality was irrelevant. Each block (50-150 trials) was preceded by a priming block (30 trials) where animals made perceptual decisions based only on a single modality presented. They then needed to continue to make sensory decisions based on that modality during the subsequent cross-modal decision-making block of trials (Fig. 1A). Our paradigm design avoided explicit per-trial contextual cues that could identify the relevant stimulus modality. Therefore mice had to learn stimuli, action, and reward contingencies together as a set to be able to infer context. Trials could be divided into congruent and incongruent trials. During congruent trials both stimuli signalled the same decision. These trials were therefore not informative as to whether the animal was attending to the correct stimulus. During incongruent trials, one stimulus modality signalled the animal to lick and the other to refrain from licking, and as such incongruent trials implied a conflict between concurrently presented stimuli. Therefore in incongruent trials, the animal had to attend to the correct modality to make the correct decision.

**Figure 1.**
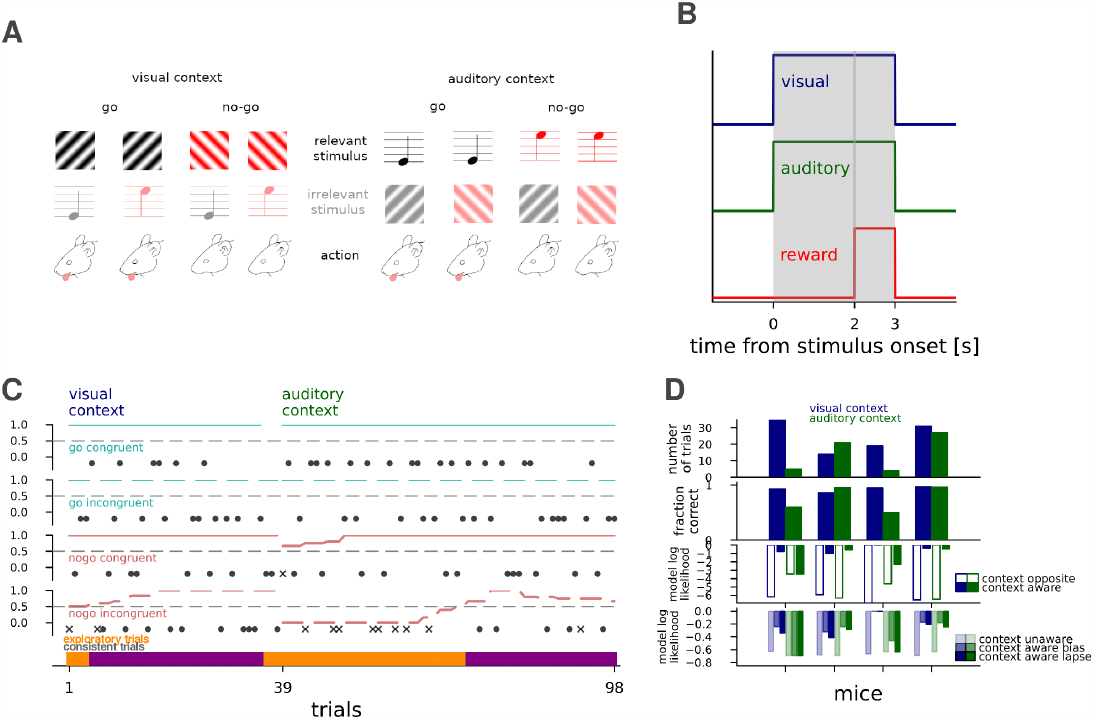
Set-shifting paradigm in mice and RNNs. **A**, Stimulus and reward structure in the set-shifting paradigm. Visual (gratings) and auditory (pure tones) are presented concurrently to mice and animals. Only one of the modalities is relevant for obtaining water reward. One of the stimuli from the relevant modality (top row) was designated as a ‘go’ signal, to which animals are expected to lick for water reward, while they were expected to withhold locking for ‘no-go’ stimulus. **B**, Time course of a trial, with simultaneous visual (blue) and auditory (green) stimulus presentation, and reward available from 2 s until stimulus end, represented as a pulse. **C**, Behaviour of an example animal in a visual to auditory context switching session for different trial types (*subpanels*). Success and failure in all four trial types are indicated by black filled circles and crosses, respectively. Lines show 21 trial equal-weight moving averages. Trials were defined as ‘task-consistent’ (*bottom panel, purple*) if the moving average performance of all four trial types were greater than chance, while other parts of the session were termed ‘exploratory’. **D**, Number of ‘consistent’ trials for individual mice with ACC recordings (n=4) for the visual (*blue*) and auditory (*green*) context (*top subpanel*). Fraction of correct responses for all trials in the ‘consistent’ periods (*top middle subpanel*, blue and green bars for the contexts as in top). Model log-likelihoods averaged over consistent incongruent trials in the visual (*blue*) and auditory (*green*) contexts for individual mice (n=4) for a model targeting the opposite modality (*empty bars*), and the correct modality (*bottom middle subpanel*). Same as bottom middle, but with a context agnostic model with mean choice lick bias (*faint colors*), and the context-aware model from the bottom-middle subpanel, but augmented with either a bias or lapse parameter respectively (*increasingly saturated colors*).

Mice were first trained on the auditory modality, then on the visual modality, and finally on the compound two-modality context switching task. The training stopped when animals showed average standard deviation from chance performance d’>1.7 (probability of chance behaviour < 0.1%). Details of the training and paradigm are described in more detail in Hajnal et al., 2021, along with the details of the behaviour of the V1 group. During each trial, visual and auditory stimuli were presented for 3 seconds with the water reward available during the third second (Fig. 1B). Intertrial interval was 3 seconds, but was extended to 9 seconds as a time-out for incorrect responses. We analysed behaviour by separating congruent and incongruent trials, and calculating a moving average for each of the ‘go’ and ‘no go’ trials separately (Fig. 1C). We defined consistent task performance as parts of the session when the moving average fraction correct of all four trial types was above 0.5. In these consistent periods, most animals performed with a high fraction of correct responses and their response statistics could be explained by context-aware models at higher likelihoods than context-unaware models (Fig. 1D for ACC and Hajnal et al., 2021 for V1, Methods). In the remaining portions they licked more randomly or with biased licking responses; these exploratory parts could feature evidence collection for context inference, reward maximization during uncertain internal task model, satiation, or fatigue (Fig. 1C bottom). 128 channel silicon probe electrodes were inserted for a single session of recording in either V1 and ACC.

We focused on delineating the neural dynamics underlying the computational steps preceding decision making in the context-switching paradigm.

### Selective suppression of irrelevant information in ACC

In the attentional set-shifting task no immediate cues are available to indicate the set of relevant stimuli. Therefore tracking the behavioral relevance of different stimuli requires that outcomes of trials are constantly monitored. We analyzed the population activity in ACC to identify the way behavioral relevance affects population responses. In particular, in the set-shifting task changes in the behavioral relevance of a set of stimuli across the visual and auditory contexts was tracked in the population activity. We constructed time-dependent linear decoders for both the visual and auditory modalities. Decoders were trained and tested on responses of ACC neurons in short time windows (50 ms) and separate decoders were constructed for each subsequent time window (Methods). Thus, the decoder could track the quality of the representation of the two stimulus modalities as each trial progressed. When decoding visual or auditory identity from ACC neuronal activity separately in the visual and auditory contexts, we found that both modalities behave similarly: irrespective of stimulus relevance, decoder performance was high following stimulus onset, but this was followed by a sharp decline in decoder performance when the modality was not relevant for making the decision. This was in remarkable contrast with decoding of visual stimulus identity in the primary visual cortex, where decoder performances did not display dependence on modality relevance (Fig. 2A,B). We interpret the consistent drop in decoder performance for irrelevant stimuli as context-dependent suppression of irrelevant information.

**Figure 2.**
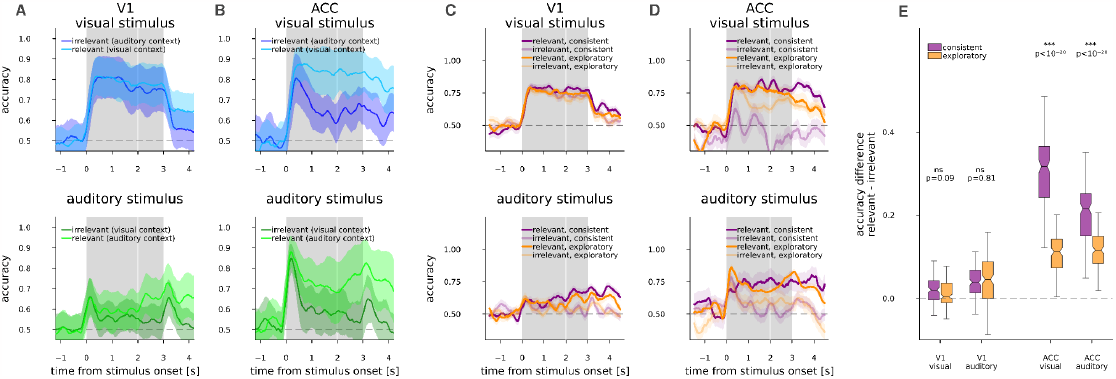
Suppression of context irrelevant stimuli. **A**, Smoothed (10 ms resolution, ma=31 points) time course along the trial of decoding accuracy from V1 neurons of visual (*top, blue*) and auditory (*bottom, green*) stimuli when they are relevant in their respective context (*light color*), or irrelevant in the opposite context (*dark colors*). Mean s.e.m. of n=8 animals, with s.e.m. also containing across animal mean of CV s.e.m. **B**, same as *A*, but decoding from ACC, n=4 animals. **C**, Similar to *A*, but trials in the relevant (*saturated colors*) and irrelevant (*faint colors*) context were stratified into consistent (*purple*) and exploratory (*orange*) groups according to the behavioural criteria detailed in Fig. 1C, while stimulus differentiating colors are omitted. Measurement in V1, n=8 animals. **D**, as *C*, but in ACC, n=4 animals. **E**, Accuracy during irrelevant trials subtracted from accuracy during relevant accuracies, the time points between 0.5-2 s from animals in *C* and *D* were concatenated and displayed as single distributions, for V1 (left four boxplots), and ACC (right four boxplots), but modalities kept separate (left and right pairs within brain areas). Significant differences in how much in irrelevant context a stimulus less decodable than in relevant context between ‘consistent’ and ‘exploratory’ trials are displayed with t-test p-value estimates and confidence levels.

Since task context and thus stimulus relevance are only inferred from contingencies between stimulus, decision, and reward, the subjective relevance of a stimulus modality might change within a task context as well, should the animal lose track of context. If the observed suppression is a consequence of the animal’s belief of the irrelevance of the given modality, we expect that in trials where suppression of the irrelevant modality is not sufficient to resolve a conflicting situation, mice would commit more errors. During a session, the behavior of mice was either characterized as ‘task-consistent’, or ‘exploratory’ in distinct blocks of trials (Fig. 1C,D). This gave us the opportunity to compare the difference between relevant and irrelevant stimulus decodability in the two behavioural states (Fig. 2C,D). Less potent suppression was apparent in ACC during ‘exploratory’ periods as opposed to ‘consistent’ periods, in contrast with V1 where no dependence on behavioral state could be observed (Fig. 2E). Behavioral state dependence of the suppression strength was consistent across the animals in ACC in both modalities (t-test, p<10^−20^ for both stimuli). No such dependence could be observed in V1 (t-test, p=0.09 and p=0.81 for visual and auditory stimulus respectively).

To summarize, context-dependent selection of the relevant stimulus is observable in mice and successful selection of the relevant modality is correlated with the success of task execution.

### Geometry of population responses in ACC

In order to understand how the observed dynamical suppression is related to executing the task, we investigated the geometry of the representation of different task-relevant quantities in ACC. Since identical stimuli are presented to the animals in the contexts and correct execution of the conflict trials require opposing decisions in the two contexts, context (whether the animal should base its decision on visual or auditory stimuli) needs to be inferred from task contingencies and represented in the activities of the neurons. We constructed time-dependent linear decoders to identify task context in population responses (Fig. 3A). Context could be reliably decoded throughout the trial but reliability varied across animals (Fig. 3A). The strength of context representation in V1 was previously shown to be stronger in behaviourally consistent periods (Hajnal et al., 2021). We investigated such a dependence in our ACC recordings as well. We did not find an improvement in context decoding in consistent trials vs. the exploratory trials (Fig. 3B).

**Figure 3.**
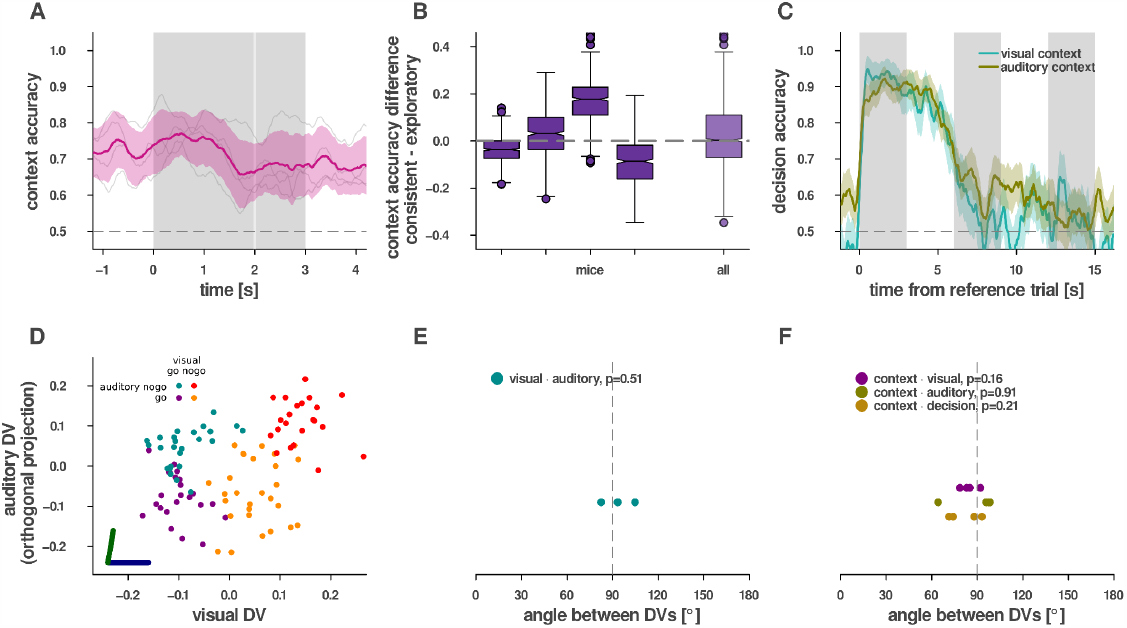
Context inference, representation geometry. **A**, CV context accuracy in ACC for individual mice (*grey faint lines*) and mean and s.e.m. over mice (n=4, *magenta*). **B**, Context accuracy difference decoded from consistent over exploratory trials for each mice individually (*left*) and the distribution for all timepoints for all mice (*right*). **C**, Decoding decision from ACC neurons of an example mouse in the reference (*leftmost grey vertical bar*), and then two subsequent trials (*all other grey vertical bars*) within the sequence for each of visual (*turquoise*) and auditory (*okker*) contexts. **D**, Population responses averaged over the first 0.5 s of stimulus presentation projected on the DV subspace in individual trials (dots), where colors correspond to visual and auditory stimulus pairing. Dark blue and green lines denote the DV directions of visual and auditory decoders, respectively. The horizontal axis is parallel to the visual DV, while the vertical axis represents the maximal projection of the auditory DV orthogonal to the visual DV. **E**, Angle between the visual and auditory DVs in ACC for each animal (*cyan dots*, n=4), mean of the first 0.5 s of stimulus presentation, p-value of t-test for mean over mice differing from 90°. **F**, Same as *E*, but between context and visual, auditory, or decision respectively (*purple, olive, light brown*).

To track the context successfully, the animal needs to keep track of the history of trial outcomes. ACC had previously been associated with representing the history of recent outcomes across trials when prediction error was high (Seo & Lee, 2007). We asked if ACC populations could decode the choice of the animal (lick vs. refrain from licking) across multiple trials. We used ACC population data to construct a time-dependent decoder for the choice the animal is making. To check the representation of a choice across subsequent trials, time-dependent decoder was not constrained to the trial in which the animal made the choice but was extended into the inter-trial interval and upcoming trials too. We found that ACC maintained information about these choices both during the intertrial interval and early in the upcoming trial too (Fig. 3C).

To understand how the neuron population concurrently represents the context variable along with stimuli of different modalities, we investigated the relationship of the representations by comparing the geometrical properties of the decoders. The decoder is defined by a hyperplane that divides the population activities in response to different values of the decoded variable (identity of context, or that of stimuli). Alternatively, the decoder can be characterized by the decision vector (DV), a vector orthogonal to the plane, which separates the identity of the decoded variable the best. The two decision vectors calculated for the auditory and visual modalities can be used to establish a two-dimensional subspace to contrast representations. Projecting neural response intensities (mean over the first 500 ms after stimulus onset) measured in individual trials into this two-dimensional subspace reveals that the auditory and visual stimulus induced changes in population activity span close to independent subspaces, highlighted by the close to orthogonal setting of the decision vectors (Fig. 3D). Such orthogonal representations ensure that different signals can be maintained without interference. This relationship between the visual and auditory decision vectors was consistent across animals (Fig. 3E, n=4, t-test, mean compared to 90 degrees, p value 0.51). We found similar orthogonal relationship between the context and the sensory or decision variables, which also held across animals (Fig. 3F, n=4, t-test, mean compared to 90 degrees, p values were 0.16, 0.91, 0.21 for context to visual, to auditory and to decision respectively).

In summary task variables are all represented in orthogonal subspaces.

### Set-shifting task in a recurrent neural network

Yet, the critical question remained as to how such orthogonal representations support set shifting. To gain further understanding of the computations required to solve the set-shifting task, we developed a recurrent neural network (RNN) computational model that mimicked the stimuli-action-reward dynamics of mice performing the task (Fig. 1B). The input to the hidden layer consisted of stimuli in the current trial as well as a variable that tracked whether the previous trial was rewarded or punished (Fig. 4A). This information has previously been shown to be represented in the prefrontal cortex (Seo & Lee, 2007) and is present in our own experimental data outlined above. We optimized the network dynamics so that it was detailed enough to explore sequential computations and memory, but coarse enough that oscillations did not typically arise at the end of the training. 30 hidden neurons yielded robust capacity for memory and computation, with input and output variables encoded in neuron pairs (one-hot encoding). Input variables consisted of the following: 1, visual input; 2, auditory input; 3, reward or punishment for correct or incorrect choices, respectively. The RNN only had hidden weights and recurrent connectivity, but no gates. A time resolution of a nine-timestep long stimulus presentation was chosen with three pre- and three post-stimulus steps without timeout on errors. The network had to decide on the response during the stimulus presentation at time step seven, and after the seventh time step reward or punishment was delivered until the next trial’s stimulus was presented. A single data point consisted of five subsequent trials from the same context in a sequence: four trials as a batch of trials during which the model had the opportunity to infer context, and a fifth final trial where the loss function was calculated from the last decision and fed to gradients for backpropagation. Training data was generated combinatorically for two choices by two modalities in 5-trial sequences, reaching 1024 unique sequences total with all possible trial combinations per context (Methods).

**Figure 4.**
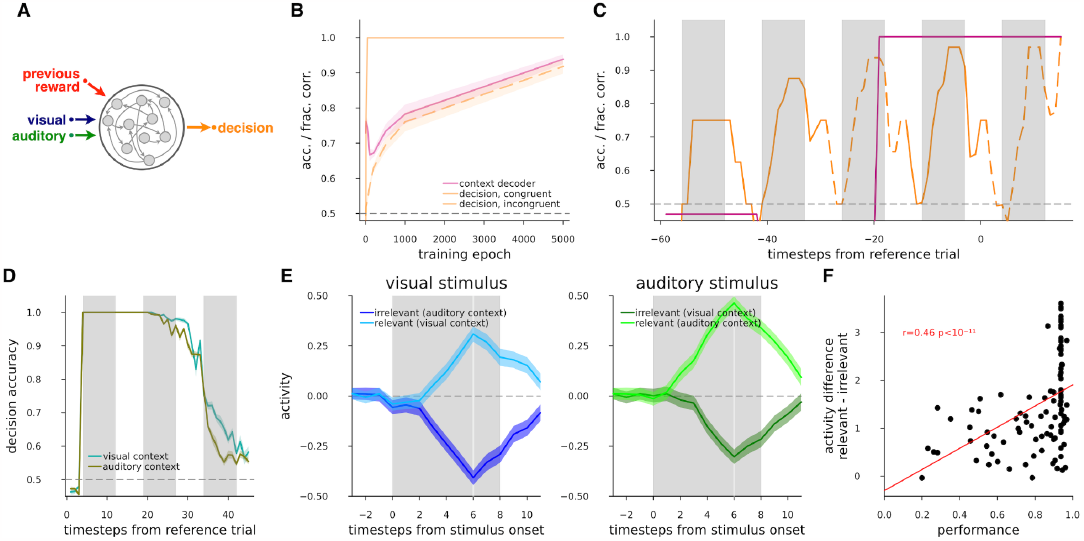
Set-shifting task in RNNs. **A**, Schematics of the RNN model **h**_t_ = f(a_t_, v_t_, r_t-1_, **h**_t-1_) and d_t_ = f(**h**_t_) at the time-resolution of trials (index t), where the decision is a map to the decision space of ‘go’ and ‘no-go’, and is a function of the hidden state vector. The update to the state vector depends on the current visual and auditory stimuli, the previous hidden state, and the reward. Stimuli and reward were switched on and off at a fine time-resolution with pulse envelopes as described in panel *C*. Note, that reward during and after a trial depended on the preceding decision, which resulted in that prior to any decision made only the previous trial was presented (hence the label ‘previous reward’). **B**, RNN performance (fraction correct), and quality of the learned representation (context decoder accuracy) as the training of the RNN progresses (epochs). Average (*lines*) and s.e.m. (*bands*) of 20 RNNs for context accuracy decoded from hidden units (*magenta*), and fraction of correct responses (*orange*) in congruent (*solid line*) and incongruent (*dashed line*) trials over learning epochs. **C**, Evolution of the decision signal and the task context when a trained RNN is performing a sequence of five trials. Decision fraction correct (*orange*) and context accuracy (*magenta*) from an example RNN after learning, for sequences with an example congruence pattern (c-c-i-c-i, where c congruent and i incongruent trials indicated by *solid* and *dashed lines* respectively as in *B*). Vertical grey bars indicate stimulus presentations. **D**, Decoding decision from hidden units of an example RNN in the reference (*leftmost grey vertical bar*), and then two subsequent trials (*all other grey vertical bars*) within the sequence for each of visual (*turquoise*) and auditory(*okker*) contexts. **E**, visual (*left, blue*) and auditory (*right, green*) stimuli when they are relevant in their respective context (*light color*), or irrelevant in the opposite context (*dark colors*), responses of the strongest n=2 (‘go’ and ‘nogo’) hidden neurons projecting to outputs combined, mean and s.e.m. from n=100 RNN models. **F**, Regression between performance of a model on all trial types (*horizontal axis*) and difference between activities in relevant and irrelevant context for all stimuli (*vertical axis*) combined from *E* for n=100 RNN models.

The task design implies that maintenance of a context variable is necessary for successful task execution. Assuming that once the correct modality has been selected the responses will all be correct, we expect that the proportion of trials where context is identified will be the same as the proportion of trials with correct decisions. Indeed, decoding context from the hidden units of RNN models, we found that the fraction of correct choices in incongruent trials were the same as the fraction of trials where the context was decoded correctly. Furthermore, after a typical performance drop in early training (epoch<200) due to strongly attuning to one of the modalities while performing poorly in the other modality, context representation and choice gradually improved in parallel, as performance became more and more context-symmetric (Fig. 4B).

From normative considerations congruent trials, where both modalities would indicate the same action, the animal has no need to rely on context information. Making a decision based on any of the two stimuli would lead to the same ‘go’ or ‘no-go’ choice. A non-trivial consequence of this is that no stimulus-action-reward contingency information could be used in congruent trials about the context, as the instructions from two modalities are indistinguishable. In contrast, indifferent to whether the decision of the animal is correct or incorrect, an incongruent trial always gives information about the current context: The reward, or lack thereof can be compared with the stimulus and response. We can observe this phenomenon in the RNN models, where a large number of sequences are available with a fixed order of trial congruence (Fig. 4C). Note, that in our RNN data sequence setup, one sequence type of four congruent and one final incongruent trials cannot be solved.

Our electrophysiological recordings in mice showed extended maintenance of the choice of the animal beyond the termination of a trial (Fig. 3C). Training a linear decoder for the choice of the RNN in a given trial we also found prolonged representation of this choice across the intertrial interval and into the upcoming trials (Fig. 4D).

We investigated the dynamics of the trained RNNs to see how selection of the relevant stimulus modality is achieved. Similar to the ACC recordings, RNNs showed maintenance of the relevant stimulus throughout the trial and suppression of the irrelevant stimulus set. Stimulus decoders cannot directly be applied to all the hidden units in our shallow RNN because they would selectively pick activity from cells with strong stimulus input. Nevertheless, we can probe the effective output of the hidden layer, by assessing the output projection weights of the hidden units to imply their importance within the hidden layer to make the decision. We found that for a hidden cell, the stronger the weight mapping to the output cells, the more it behaves as a modality-invariant abstract cell. Thus, it responds to the context-relevant stimulus input with enhanced response, and to the irrelevant stimulus input by a negative response (Fig. 4E). Note, that our RNN model did not have any preference for the sign and magnitudes of weights and activities except through weight regularization; hence suppression of irrelevant and enhancement of relevant activity were symmetric.

In summary, our RNN model reproduces the performance and neural activity of mice in a set-shifting task, such as the geometry of the representations of the stimuli and the context. The model learned to infer the context from congruent and incongruent trials, and to suppress the irrelevant stimulus modality.

### Context-gated attention in activity subspaces

Having reproduced some of the characteristic behavior of the recorded ACC neuron populations, we surmised that the RNN could be a promising tool to investigate the question of how these observed representations may support set shifting. For this, we first considered the attentional selection problem, then we identified a potential computational mechanism that can drive attentional selection, and finally we derived experimentally verifiable predictions.

Selection by attention can be regarded as a way to perform cognitive abstraction. The selection of the relevant quantity is not trivial since i) dimensional reduction needs to occur to map the space of all possible stimuli with various combinations of simultaneous, potentially conflicting instructions to a single dimensional final action space, and ii) this mapping depends on the context. To simplify the argument, in this section we assume that the information about the context is already available in the animal. We also assume that the task is learned, and connection weights do not change.

In the space of population activity, we describe the activity evoked by the visual and auditory stimuli by vectors **v** and **a**, respectively. Thus, the activity arising as a consequence of joint presentation of the two stimuli is simply **v** + **a**, which resides in a space that is a direct sum of the orthogonal subspaces spanned by the visual and auditory activity we observed. This combined space is a subspace of the complete neural activity space (Fig. 5A). The animal is required to compute a decision based on these stimuli. The decision will be characterized by an activity vector **d**. The transformation of the activity from sensory stimuli to the decision can therefore be described by the transformation **d** = F(**v** + **a**): F projects the stimulus subspace to the smaller decision subspace. In incongruent trials the animal is expected to learn to arrive at different decisions for identical stimuli such that the transformation F that is implemented by a network of neurons is invariant across contexts. Therefore contextual selection can only work if the stimulus subspace has additional modulating activity. We describe this modulatory activity as an additional component in the activity space, denoted by **m**. Thus, considering an additive modulatory signal decision is actually acting on the joint activities d = F(**v** + **a** + **m**) (Fig. 5A). In order to remove dependency on the irrelevant modality the modulatory signal shall either i) remove that modality, e.g. in the visual context elimination of **a** requires **m** = -**a** (Fig. 5A), or ii) can also enhance the relevant subspace: **m** ∝ **v** in visual context and **m** ∝ **a** in auditory context. A context dependent modulatory signal **m** can thus be formulated as applying mapping **M** to context **c**: **m**(**c**) = **Mc**. Intuitively, (i) suggests that **m** is suppressing **a** in the visual context and vice versa in the other context. Critically, the only transformation that fulfils this requirement is mutual inhibition of the two stimulus subspaces, corresponding to a matrix with a qualitative structure of 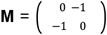 (Fig. 5A). We provide formal proof in Proposition 1 and Corollary 1 (Appendix). Less formally, we argue that subspaces of neuronal activity that are selective to a particular stimulus modality will display strong modulation with changes in context. A scaling of **M** corresponds to imperfect suppression of the irrelevant modality. Note, that from the point of view of the abstract decision, enhancement of the relevant stimulus modality is equivalent to suppression (Appendix). Such enhancement can be achieved by having positive diagonal elements in **M**. Enhancement and inhibition acting together, each with the strength of the matrix elements yields the classic picture of a neuron pair with mutual feedback (Fig. 5B). We generalized these statements in Corollary 2 (Appendix) to include mixed cell selectivity with rotation, interfering modulation on non-orthogonal stimulus subspaces, while we also extended linear subspace representations with biologically plausible nonlinearities and sequential processing. Most importantly, we conclude that when connection weights are fixed, the computation of context-dependent selection of the relevant stimulus has to act on the stimulus subspaces, enhancing the relevant stimulus and inhibiting the irrelevant subspace.

**Figure 5.**
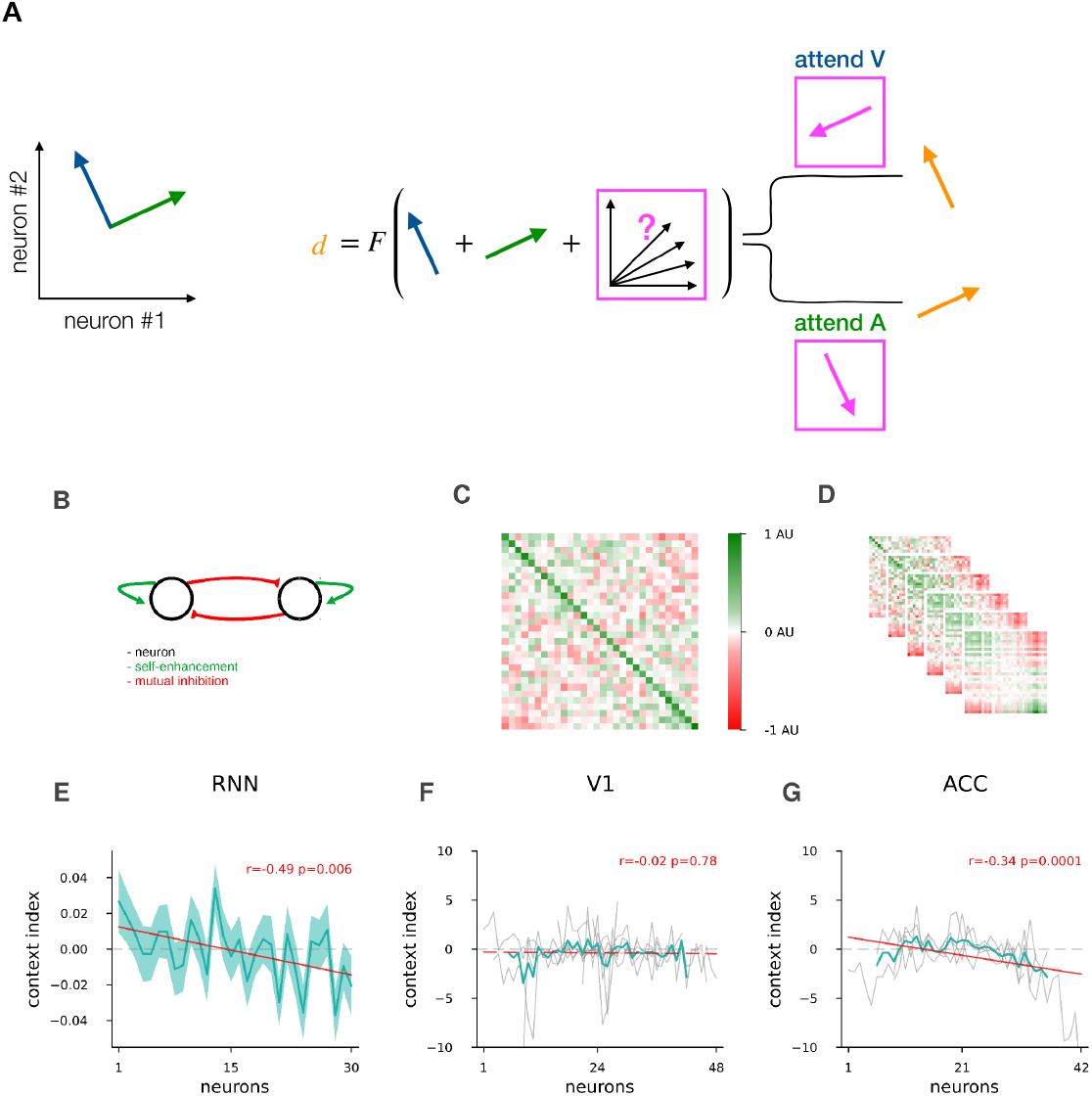
Context gated mutual feedback mechanism. **A**, Schematics of a context-selective inhibition mechanism when representations can be approximated by linear subspaces. At the level of primary sensory areas, the total stimulus space consists of unrestrained activities along each subspace (*blue* and *green, left*). In order to map the total stimulus space to the abstract decision space (*orange, right*) in a contextually correct way with fixed weights, the activity in the irrelevant subspace has to be inhibited (*pink, middle*). **B**, Schematics of mutual feedback in the total stimulus subspace represented by an axis aligned two dimensional system: two neurons with mutual inhibition and positive self-enhacement. **C**, Recurrent weight matrix of RNN models; neurons ordered by their modality index defined as input weight difference between visual and auditory stimuli. Colormap rescaled by the largest of absolute minimum or maximum (AU). Neurons ordered separately for ‘go’ and ‘no go’ signals, then averaged, mean of n=100 models. **D**, same as *C*, but also the 2^nd^ to 6^th^ power of the recurrent weight matrix on *C*, approximating its final recurrent effect from stimulus onset until decision requirement, colormap rescaled, units as in *C*. **E**, Context index defined as difference of neural activity between the visual and auditory context; neurons ordered by the modality index, for RNN models, using input weight modality index as in *C,D*, mean and s.e.m. over models, n=88 that have performance > 0.9 fraction correct. Values calculated and neurons ordered separately for ‘go’ and ‘no go’ signals, then averaged. **F**, same as *E*, but mouse modality index was defined as trial-averaged response difference between visual and auditory stimulus, when presented in the absence of the other stimulus. Mice had varying numbers of neurons, with the neuron index set at the midpoint of the number of neurons for display purposes. In V1, Individual mice (n=8, *grey lines*), mean over mice skipping 4 start and end neuron order positions on each side (*turquoise*), and linear fit to mean over mice (*red*). **G**, same as *F*, but for mice recorded at the ACC site, n=4.

The above argument indicates that context-gated suppression is a potent mechanism to perform the set-shifting task. We next investigated if this mechanism corresponds to the computations that are implemented in our RNN. In our RNN, the recurrent connections, F, has to perform all the computations necessary to reach a decision from the auditory and visual inputs: since context **c** is represented in the RNN, it is transformed by recurrent dynamics into modulation **m**, which in turn is integrated with the stimulus representation **v** + **a**. First, we examined if the trained RNN displays the mutual inhibition pattern required for suppression of the irrelevant modality. We investigated the relationship between the input connections to and the recurrent connections within the hidden layer in RNNs. We therefore calculated a modality index by determining the difference between the strength of the input weights conveying the visual and auditory inputs for each hidden unit. We sorted the hidden neurons by the modality index, so that the first neurons receive strong visual and weak auditory input, while the last neurons weak visual and strong auditory input. We investigated the network structure by displaying the connection matrix for the recurrent layer of the RNN. The connection matrix is largely unstructured but revealed patterns reminiscent of the transformation F: diagonal elements showed positive feedback, and far ends of the modality index spectrum revealed traces of inhibition (Fig. 5C). One can more clearly visualize the transformation performed by the RNN by unfolding the dynamics of the RNN across the trial. Stimulus onset and decision making were separated by six time steps. To reveal how inputs are transformed by the RNN by the time decisions are made, we plotted the 6th power of the recurrent connection matrix, linearly simulating repeated multiplication on the hidden states. We observed a strengthening of the positive self-feedback and the mutual inhibitions onto disjoint block matrices (Fig. 5D). Taken the results for suppression and inhibition together, the structure of the lateral connections is characterized by a simple block matrix structure of 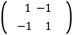.

Second, we sought to establish a signature of inhibition-based manipulation of stimulus representations. The mutual inhibition between the two stimulus modality-specific neuron groups causes each group to be less active in their irrelevant context after suppression. Hence the firing rate differences between activities in the two contexts, i.e. the context representation, will at least partly originate from the context-gated suppression itself. The two computations, the slower context inference and the faster irrelevant suppression, are tied together during active attentional modulation. This coupling can be tested in the trained RNN by establishing the relationship between the modality specificity and contribution to context decoding of individual neurons. We constructed a context index for individual neurons by taking the difference in the activity of each cell between the two contexts. To ensure that the stimulus statistics are the same in the two contexts we only used congruent trials. We found significant correlation between the context index and the rank of the neurons when the neurons of the hidden layer of the RNN were sorted according to the modality index (Fig. 5E, r=-0.49, p=0.006).

We tested the prediction that neurons displaying higher modulation by stimulus modality changes will be more intensely modulated by changes in context in recordings from mice. To explore the specificity of the prediction to ACC, we analyzed neurons recorded in both ACC and V1. For the modality index we calculated the difference between average responses to visual stimulus and to auditory stimulus. In order to compare neuron responses to each stimuli without interference from the other stimulus modality, we used single modality trials, not part of the complex task, but recorded in the same session. Similar to the RNN analysis, we sorted the neurons according to their modality index. As cells typically encoded either the ‘go’ or the ‘no go’ signal, modality and context indices were calculated and sorted separately for the two signals, and then combined. The context index was calculated for individual neurons based on the difference of responses in the two task contexts. We found a clear correlation between the context index and neuron modality rank in mice in ACC (r=-0.34, p=0.006, Fig. 5G), but not in V1 (r=-0.02, p=0.78, different between areas, z-test p=0.004, Fig. 5F). Therefore neurons in ACC that are selectively engaged in one of the stimulus modalities tend to display strong modulation with task context.

In summary, identifying subspaces of stimuli and context within both recorded ACC neurons and the hidden layer of the RNN model revealed the same gated mutual feedback we proved theoretically. The effect of this mutual feedback apparently results in a particular correlated structure of cell responses: Cells that respond more selectively to a stimulus modality tend to be more active in the context where that modality is relevant. This essentially means that the representation of the attention cue, context, and the selection target, modality, is largely overlapping in the same neural population. In this configuration, downstream decision making centers can dynamically read out the abstracted relevant stimulus representation from the total stimulus subspace without changing the readout weights.

## Discussion

We examined the computations required to establish the relevance of incoming stimuli and the way this computation is reflected in the activity of neurons recorded in ACC. In a changing environment where relevance of stimuli can change, we argue that resolving conflicts between concurrently presented stimuli is key to effective performance. We found that suppression of irrelevant stimuli was a signature of conflict resolution in ACC as suppression prevalently occurred when behavior also reflected ignorance towards the irrelevant stimulus. We identified analogous suppression in an RNN model of the task. We proposed that context-based attention is a critical mechanism for dynamically identifying relevant stimuli from a wider set and we found that the geometry of the representation in the ACC efficiently supports context-gated attention. In addition, we confirmed a conjecture of theoretical considerations obtained by analytical derivation that recurrent weights of the trained RNN models achieve enhancement of current context and suppression of irrelevant context through mutual inhibition. The RNN that was trained on the set-shifting task predicted that context representation and the representation of the two stimulus modalities are dependent, which prediction we confirmed in our population recordings from ACC. Neither suppression of the distractor, nor the correlation between modality specific and context specific activity was observed in V1, suggesting that these computations are performed in higher areas, a signature of which we identified in ACC. Thus a context-gated attention mechanism can dynamically resolve instruction conflict during presence of a distractor.

Our recurrent network model provides important insights into the way relevant stimuli can be identified in a rich environment through the contribution of population responses to solving the set-shifting task. The mutual suppression structure of the lateral weights in the set-shifting task-trained RNN supports the amplification of the currently active modality over the currently inactive modality. This architecture in turn supports the maintenance of the current context. Thus, the synaptic weights of the network have a dual role: they both contribute to the modulation of the total stimulus subspace, i.e. the modality targeted by attention, but also contribute to the maintenance of context. This dual role helps understanding the relationship we demonstrated between the context and modality indices of individual neurons.

The activity representing the contextual cue and the target modality of the attentional selection mechanisms are highly correlated; a prominent pattern that emerged from our results. We argue that this correlation is exactly due to the suppression of the irrelevant modality. We showed that the cells responsible for the representation of the non-suppressed modality are the ones generally more active during that context. Note, that the trained RNN is both capable of maintaining a constant representation of the incoming stimuli and of representing them in a context-gated manner. This is also characteristic of the representation identified in ACC as evidenced by unaltered activity in the first 500-ms of stimulus presentation and context-driven suppression later in the trial. Similar geometry was proposed for attentional selection in Mante et al., 2013 as a context-dependent stimulus-selector vector. Here we proved that this modulation is the only possible computational mechanism to shift attention dynamically. Furthermore, we demonstrated that this computation emerges through specific mutually inhibiting input and lateral connections

The computational motif of mutual inhibition between subspaces representing different modalities was identified i) analytically, ii) in RNN models, and iii) in ACC of behaving mice. The analytic derivation and the recurrent feedback connections of the RNN model explicitly revealed an architecture characterized by mutual inhibition. Modelling studies showed that this connectivity allows switching between distinct dynamical behaviors (Machens et al., 2005). Although connection structure between concrete neurons was not available with typical electrophysiological recordings, anatomy of the local circuitry in the cortex still provides insights into the relevant circuitry in ACC. Multiple inhibitory neuron types have been identified that are coupled to pyramidal neurons that control local circuit output and can potentially contribute to lateral inhibition (Karnani et al., 2016). Such lateral connections are typically part of competing representations in sensory cortices (Machens et al., 2005; Vandrey et al., 2022), in the prefrontal cortex (Strait et al., 2014), and in the brainstem (Koyama & Pujala, 2018). Although no such studies are available for ACC, it is likely that similar anatomical motifs contribute to both stabilizing context representation, and upon prediction errors computing the correction of a mismatched context representation in the ACC.

In the ACC, task variables, e.g. stimuli and context, jointly modulated the activities of individual neurons, resulting in a mixed-selectivity representation. Mixed selectivity has been demonstrated in multiple brain areas (Hajnal et al., 2021; Mehta et al., 2019; Parthasarathy et al., 2017). Traditionally, it is the specific selectivity of neurons to certain variables that has been identified (Hubel & Wiesel, 1959; O’Keefe et al., 1978; Solstad et al., 2008). While the observed phenomena, such as selective suppression, could be sought in individual neurons, we focussed our analyses on population responses. Our analyses, which identify low-dimensional subspaces that accommodate orthogonal representations for task variables, provides an important insight into the relationship between mixed and selective representations: in a rotated coordinate system activity along a particular axis covaries with a single independent variable. As such activity of a mixed-selectivity neuron belongs to multiple rotated linear subspaces, while a specific selectivity neuron can be considered to belong to an axis-aligned subspace. Accordingly, while linear readout in this mixed-selectivity space is a potent computational tool, readout weights need to take into account the rotated representation (Rigotti et al., 2013; Ruff & Cohen, 2019).

In order to investigate how changed relevance of sensory stimuli is processed in the cortex, we developed a paradigm where the context was not presented to the animals but had to be inferred through trial and error. If the context is cued there is no actual conflict after learning the task and context-gating function is attributed to the PFC (Mante et al., 2013). Beyond conflict resolution, in our experiments ACC might be implicated in reinforcement learning related computations (Holroyd & Coles, 2002): the performance of our animals saturated below 100% correct responses as they made a number of exploratory trials to optimize their behavior (Fig. 1C). We conclude that when a task is not fully learned and the animal is frequently faced with situations where conflicts arise, the ACC plays a larger role preparing the PFC for the optimal task representation, while post-practice decrease in ACC activity is larger than in the PFC (Khamassi et al., 2015; Milham et al., 2003).

Evidence from our mathematical analysis indicates that unless the animals actually performed this inference they could not successfully perform the task. Earlier research has indicated that components necessary for context inference are represented in the cortical circuitry. PPC has been shown to maintain a history of past environmental information (Akrami et al., 2018), while PFC has been implicated in maintaining previous choices and rewards (Seo & Lee, 2007). Interestingly, the contextual representation appears to be continuous in ACC and V1 (Hajnal et al., 2021), as well as in our RNN model: the context signal is present during off-stimulus periods. This indicates that the context representation is maintained at a time scale beyond that of a single trial. This is consistent with the idea that storing information in short term memory by vmPFC (Moneta et al., 2021) is more effective than recomputing a constant context on each trial. Nevertheless, the representation should be flexible so that it can be updated if context changes. ACC might be involved in contextualizing goal-oriented sets of stimuli, action and reward upon encountering error feedback (Shen et al., 2015). We did measure context-related activity in ACC of mice, thus it could potentially be used to resolve conflict by ACC. However, whether computations required to infer context is mainly computed locally in ACC or a larger fraction of context-related activity originates from other brain areas could not be uniquely determined by our recordings.

We used decoders to identify context-related signals in ACC. Note, that decoders pick up any source of neural activity pattern that differs between the two blocks of a session. Therefore it is not entirely surprising that there appeared to be a contextual signal even when the animals’ behavior did not reflect proper task execution, both shown as error trials, and as longer exploratory periods. The signal identified by the context decoder under such unsuccessful conditions might come from different sources: i) an uncertain representation of context that demands more sampling of the task space, ii) environmental non-goal-oriented context information, which may come predominantly from PPC (Akrami et al., 2018), and iii) an asymmetrical task execution, in which one of the contexts is well successfully performed but in the other context the absence of goal-oriented knowledge introduces response modulations. We hypothesize that during unsuccessful task execution the outcome history contingencies and prediction errors in ACC can both contribute to a non-goal-oriented context representation (Brown & Braver, 2005).

We emphasize that correct execution of the uncued set-shifting task relies on a number of computational steps to be executed, namely context inference and selection of the relevant set. Disruption of any of these seems to result in declining behavioural performance for the animals, with specific signatures that are identifiable in our model as well. Incorrect context inference causes the animal to switch to alternative strategies that are most evident in incongruent trials as: i) randomly choosing between licking or withholding lick; ii) reclining to a lick-only strategy; or iii) making choices according to the rules of the other context, i.e. making decisions based on the distracting stimulus. Our RNN model does not implement complex strategy-exploring algorithms, however strategy “iii” observed in mice was clearly identifiable in the RNN model in the early epochs during learning: In one of the contexts, incongruent trials elicited choices that would have been correct only in the other context. Interestingly, our analyses revealed that mice performed worse in trials where the suppression of irrelevant subspace was not sufficiently strong in ACC. Our model with an emergent simple mutual inhibition reproduces context-specific suppression in a well trained model. Accordingly, in early learning epochs a strategy reminiscent of “iii” was coinciding with a lack of the suppression of the distractor modality.

The aim of the context-gated attention is to prepare the correct set of stimuli for a downstream target of ACC, the PFC (Spellman et al., 2021), by selecting the relevant stimulus. Our results demonstrate that in attention-shifting tasks this is achieved by constraining the activity of ACC to a smaller subspace that only accommodates the relevant modality. This computational mechanism has the appeal that a downstream area can read out information relevant for a choice relying on fixed connections that project from the entire stimulus related subspace (Bernardi et al., 2020; Murray et al., 2017; Ruff & Cohen, 2019), thus resulting in an efficient solution for conflict resolution.

## Appendix

### Proposition 1

Assuming linear neural activity space representation, with fixed weights, modulatory signals must act on subspaces.

*Proof*. To formalize the problem, suppose *X* is a vectorspace that represents the space of possible neural population activity. The coordinates for **x** ∈ *X*, x_1_,x_2_,…,x_N_ are instantaneous firing rates of each of N neurons. Task related variables can be represented in subspaces of *X*. The results below are equally applicable to more than one dimensional stimulus representations, including two dimensional dependent one-hot encoded otherwise one dimensional stimuli, as well as to multiple modulations: In the equations below only the bases need to be expanded, connection matrices changed to block matrices, standard basis vectors changed to the sum of multiple standard basis vectors. For simplicity of notation, here we restrict the proof to two stimulus subspaces, and minimal encodings in one-dimensional linear subspaces. In addition, as per the premise above, we are dealing with agents that already learned the task, so we will not explore dynamically changing weights. Stimulus encoding subspaces can be found by linear decoders projecting with coefficients **e**_V_, **e**_A_ ∈ *X* to *V* and *A*, the subspaces of *X* where the visual and auditory stimulus related activity resides. We use the format, **e**_S._, as basis for one dimensional subspaces *S* ⊆ *X* expressed in the native coordinate system of *X*. Note, that *V* and *A* are subspaces, as long as linear independence holds, i.e. while the absolute value of the correlation between visual and auditory related activity < 1, but this is not enough for non-interfering modulations. We focus our assessment on orthogonal subspaces (where correlation is 0, and **e**_V_ · **e**_A_ = 0), and expand on linearly independent non-orthogonal subspaces in Corollary 2 (iii). We will also assume stimulus input to the neurons in *X* are constant, *v* and *a*, so any further manipulation to stimulus has to happen within *X*. In this setup the total stimulus related activity, **s**, (Fig. 5A) is:

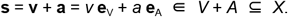

The task requires that *X* has an abstract one dimensional subspace, *D* ⊆ *X*, which has activity **d** = *d* **e**_D_ ∈ *D* that represents the final output, i.e. the ‘decision’ of these neurons. The map to this decision subspace can be performed by a linear operator F ∈ **ℒ**(*X*). With the synaptic weights as components of the map, and initial activity **x** ∈ *X*, the transformation is

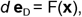

or in neural coordinates:

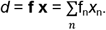

At this point we introduce a modulation *u*, that changes *d*. Both the map F, or the activity, **x** can depend on the modulation, but the weights f_n_ are constants by the conditions of the proposition. This leaves us to implement modulation dependent computation within **x**. We define stimulus-unrelated activity, **z** ∈ *Z*, in the complement of *V* + *A*, so that (*V* + *A*) ⊕ *Z* = *X*; **z** can also have modulation-dependent terms, hence **z**(u). Thus we need to solve the following problem for a linear modulation map, **m**(u) ∈ *X*, that maps the modulation, *u*, to *X*:

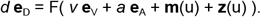

First, it is clear that the dot product **m**·(**e**_V_+**e**_A_) ≠ 0, otherwise **m** does not influence stimulus related activity. Second, by definition we have incorporated into **z** any stimulus-unrelated terms from the basis decomposition of **m**(u), including modulation-dependent stimulus-unrelated terms, denoted as **z**(u). These two items can only occur simultanously, if **m**(u) maps only to the subspace spanned by the linear combination of **e**_V_ and **e**_A_., i.e. to *V* + *A*.

Thus a modulating signal that aims to influence the decision which in turn depends on stimulus related activity must map onto the stimulus subspace.

## Definition

Context is defined as the set of stimulus the decisions need to be based on. Thus in each context one set is relevant and the others are irrelevant. More specifically in our paradigm the contexts are defined by the relevant stimulus modality.

## Corollary 1

Context modulation gates activity on the stimulus subspaces by self-enhancement and mutual inhibition.

*Proof*. Let us formalise context modulation. Let the modulation map **m** from Proposition 1 depend on a vector valued context modulation term, u = **c**. In *X* coordinates

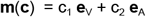

where 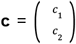 is the one-hot encoded context vector that takes values 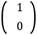 or 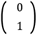 for visual or auditory context, respectively. It is clear that for self-enhancement, the coordinates of c are in the appropriate order, while for inhibition the equation has the form of **m**(**c**) = - c_2_ **e**_V_ - c_1_ **e**_A_. This notation has some complicated case by case description requirements, so we improve on the notation.

We would like to express **m**(**c**) in matrix notation; the benefits of why it is a useful notation will be expanded below. We start with the case for inhibition. Observe that **m**(**c**) has to contain a multiplicator, 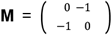, that is anti-diagonal, signifying that the irrelevant subspace is the opposite of the context-relevant subspace corresponding to the changed order and negative sign above, and with negative components so that its effect is inhibition:

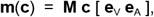

Where [.] is a matrix from column vectors for the basis vectors of *V* + *A*. Applying this context dependent modulation as inhibition to the stimulus related input yields the correct order and sign as above in the simple notation:

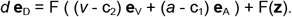

The most efficient inhibition would reduce the activity in the irrelevant subspace to 0. This can be achieved with 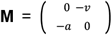. Although the decision can be further influenced by F(**z**), it cannot have stimulus-related projection, as *Z* and *V* + *A* are disjoint by definition. Thus **m**(**c**) contains all that is both allowed and needed for context specific inhibition of irrelevant stimuli.

The above matrix multiplication in **m**(**c**) gives an intuitive implementation in neural circuits: Context, one-hot encoded in vector **c** as defined above, acts as an input to the stimulus subspace, *V* + *A*, through fixed input connections **M**, resulting in context-gated mutual inhibition. As a concrete example, the effect of this gated inhibition with F = (1 1), the simplest additive projection mapping to *D*, while disregarding **z** for simplicity, expressed in *V* + *A* coordinates, in the visual context, is:

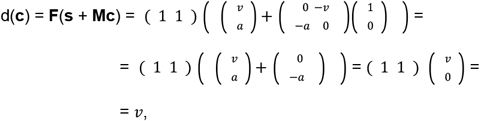

thus only the visual activity is projected onto the decision subspace. Note, that we did not assume the exact place in the brain hierarchy where the inhibition should take place, in other words *X* can be arbitrarily large. It is just necessary that it happens before mapping to the final action space by F. The concrete implementation of inhibition has many biological mechanisms, but ultimately all can be transformed into the format in the proof: Reducing outgoing activity of specific neurons that form the coordinates of the basis for that stimulus encoding subspace.

An enhancement of the relevant stimulus instead of, or beside the suppression of the irrelevant stimulus is also plausible. If the modulator input connections **M** is changed to a diagonal matrix with any positive components **υ** and 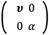, it will act as context gated positive feedback. With these connection weights the proof holds for selection with an additional condition. The downstream readout threshold (below which input is discarded) has to be set between the enhanced and normal stimulus activity levels; this condition is often true as thresholds are typical parts of neural circuitry. One can combine self-enhancement and mutual inhibition of subspaces with the qualitative form 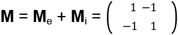.

In summary we proved that in linear subspace representations, a simple context driven switch is capable of selecting the relevant modality with inhibition or enhancement, and with fixed weights this computation has to act on the stimulus subspaces.

## Corollary 2

The proof of Corollary 1 can be generalized with regard to F. The F function on *X* can typically have five additional characteristics in neural circuits: i) it can expand out of the stimulus subspace, ii) connections can allow mixed selectivity, iii) can modulate in non-orthogonal stimulus subspaces, iv) the mapping is composed with nonlinear transfer functions, v) brain circuits and RNNs often compute in sequential activation.

*Proof*. We address these four problems separately.

i) We can simply disregard any component of F that escapes its interesting part of its range, *D*, as it will not influence downstream decision making.

ii) Although many cells typically operate with mixed selectivity, a number of variables can still be encoded simultaneously within a set of neurons without interference: The basis vectors of such mixed encoding subspaces are rotated from the natural neural coordinate system, preserving orthogonal relations between correlated activity directions, i.e. subspaces. The above proof works in these rotated subspaces as well, **F** feedback connections and **D** decision projection can be transformed so that they operate either or both on rotated domain or range spaces. With the additional use of simple change of basis operators the form of the proof remains the same.

iii) when **v** and **a** are not orthogonal, the modulation **m** will have components that also modulate the relevant subspace to the extent of the angle between the two subspaces. Still, non-orthogonal stimulus subspaces are modulated in a way that the largest modulation is in the intended direction, while to some extent it spills over to the unintended direction. Therefore fully orthogonal subspaces are of special interest, because they have zero projection onto each other, therefore the modulation is non-interfering with irrelevant subspaces.

iv) There can be arbitrary nonlinear computations if a σ nonlinearity is applied: σ ∘ F. However if σ is smooth and monotonic, which is true for most biological neurons, the shape of the nonlinearity will not affect the local subspace geometry, nor will it affect the additive inhibition or enhancement. A notable exception is negative sign change by σ, but that can be addressed by flipped connection weights in elements of F if the operating range of σ requires it, as both F and σ are fixed. Thus all statements on subspaces are also applicable in submanifolds with smooth differential structure.

v) Sequential F can be thought of as a composition F ∘ F ∘ F ∘ … or **F**^*t*^ after time *t* before opening the gates for output. At the required time point, however, the abstraction has to confine the meaningful decision related output activity onto *D*. Should F map stimulus related activity outside D + V + A, an extension to ii), the original *V* + *A* subspace will just need to be unified with the subspace of the range of F where *V* + *A* got mapped, and the above proof works with this larger *V*^*^ + *A*^*^ ⊆ *X*, and *V* ⊆ *V*^*^, *A* ⊆ *A*^*^. A simple consequence of repeated application of F is that F can take over the role of mutual feedback of the input M for maintained activity, for example, when the input **c** is for a single time point, that gets mapped onto *V* + *A*, and then can be maintained by F with self-enhancement and mutual feedback similar to **M** for the subsequent time points within *V* + *A*. This essentially allows for memorizing the short input signal, and maintaining it even in absence of a clear task, or stimulus, but within the stimulus subspace *V* + *A*. However, without the initial context input, even with **M**-like F, the bistable system on its own will not decide on the projection and will stay on its unstable point, eventually randomly landing on one of the fixed states.

Thus, the proof for Corollary 1 also holds on more realistic nonlinear, larger dimensional, sequentially operating neural manifolds.

## Authors’ Contribution

P.O.P., M.E, and P.G. designed and optimized the behavior, and designed all experiments. D.T., M.E. and K.S. trained animals and performed the recordings. D.T, M.V.M., K.S. and M.A.H. curated the data. M.A.H. and G.O. designed the analysis. M.A.H., Z.S., A.A. performed the analysis with input from G.O. M.A.H., Z.S., A.A. developed the RNN and performed the analysis. M.A.H., Z.S., G.O. and P.G. wrote the manuscript.

## Acknowledgements

P.G. and G.O. were supported by a grant from the Human Frontiers Science Program, P.G. was supported by grants 1R01MH105427, R01NS099137, 1P50HD103557, G.O. were supported by a grant by the Hungarian Brain Research Program (2017-1.2.1-NKP-2017-00002). This research was supported in part by the National Science Foundation under Grant No. NSF PHY-1748958 (G.O.).

## Competing Interests

The authors declare no competing interests.

## Methods

## Surgery

All experimental procedures were approved by the University of California, Los Angeles Office for Animal Research Oversight and by the Chancellor’s Animal Research Committees. 7-10 weeks old male and female C57Bl6/J mice were injected intraperitoneally with analgesic (carprofen, 5 mg/kg of body weight), anesthetized (isoflurane, 3–5% induction, 1.5% maintenance). Stainless steel head holder was implanted on the skull using Vetbond (3M) and dental cement (Ortho-Jet, Lang), and a recording chamber was built. Mice recovered with carprofen (2 days) and amoxicillin (7 days) administered. Approximately 1 day before the recording, a silver chloride ground wire was implanted within a small craniotomy (0.5mm) above the right cerebellum and fixed in place with dental cement. A circular craniotomy (diameter = 1 mm) was performed above the ACC (anterior-posterior 1.8 mm, medio-lateral 0.25 mm). The exposed skull and brain were covered and sealed with a silicone elastomer sealant (Kwik-Sil, WPI).

## Animal training

Animals were first trained to perform unimodal visual and auditory lick/no-lick (go/no-go) discrimination tasks. Licks are detected by using a lickometer (Coulbourn Instruments). Lick detection, reward delivery and removal, sensory stimulation and logging of stimuli and responses were all coordinated using a custom-built behavioral apparatus driven by National Instruments data acquisition devices (NI MX-6431) controlled by custom-written Matlab code. A 40-cm (diagonal screen size) LCD monitor was placed in the visual field of the mouse at a distance of 30 cm, contralateral to the craniotomy. Visual stimuli were generated and controlled using the Psychophysics Toolbox (Brainard, 1997) in Matlab. In the visual discrimination task, drifting sine wave gratings (spatial frequency: 0.04 cycles per degree; drift speed: 2Hz; contrast: 100%) at 45 degrees, moving upwards, were paired with a water reward. Drifting gratings of the same spatial frequency but at 135 degrees orientation, moving upwards, signalled a reward would not be present, and the animal was trained to withhold licking in response to the stimulus. The inter-trial interval was 3 seconds, except for trials in which the animal had a miss or false alarm, then the inter-trial interval was increased to 6.5 seconds. The animal’s behavioral performance was scored as a d’ measure, defined as the z-score of the hit rate minus the z-score of the false alarm rate. Once animals reached expert performance (d’>1.7, p<0.001 as compared to chance performance, Monte-Carlo simulation), they were advanced to learning the auditory discrimination task where a low pure tone (5 kHz, 90 dB) indicated that the animal should lick for reward and a high tone (10 kHz, 90 dB) indicated that the animal should withhold licking. The inter-trial interval was similarly 3 seconds and the inter-trial interval was increased to 9 seconds after misses or false alarms. After animals learned the auditory discrimination task (d’>1.7) they were trained to perform the multimodal attention task. In this phase, animals first performed one block of visual discrimination (30 trials). If their performance was adequate (d’>2.0, correct rejection rate>70%, hit rate>95%) they then performed the visual discrimination task with auditory distractors present (the high or low tones) for 120 trials. Then, after a five-minute break, they performed the auditory discrimination task for 30 trials and if their performance was adequate (d’>2.0, correct rejection rate>70%, hit rate>95%, further analysis was conducted during the recorded session shown in the results), they performed auditory discrimination with visual distractors present (oriented drifting gratings at 45 or 135 degrees described previously). During each training day and during the electrophysiological recordings, each trial set started with 30 trials where only visual or auditory stimuli were delivered which signalled whether the animal should base its decisions on the later multimodal trials to visual or auditory stimuli respectively. Each trial lasted 3 seconds. When the cue stimulus instructed the animal to lick, water (2 microliters) was dispensed two seconds after stimulus onset. No water was dispensed in the no-lick condition. To determine whether the animal responded by licking or not licking, licking was only assessed in the final second of the trial (the response period). If the animal missed a reward, the reward was removed by vacuum at the end of the visual/auditory stimulus. Animals performed 300-450 trials daily. Only one training session was conducted per day with the aim to give the animal all their daily water allotment during training. If animals did not receive their full allotment of water for the day during training, animals were given supplemental water one hour following training. Whether the animal started with the attend-visual or ignore-visual trial set was randomized in every session. Importantly, the monitor was placed in the exactly the same way during the auditory discrimination task as it was placed during the visual discrimination task, and a grey screen, which was identical to that during the inter-trial interval of the visual discrimination task and isoluminant to the drifting visual cues, was displayed throughout auditory discrimination trials. As a result, the luminance conditions were identical during visual and auditory discrimination trials.

## Behavioral analysis

Performance of the animals was characterized by 20 trials wide sliding window average. Four components were constructed for ‘go’ and ‘no-go’ signals, each having two for congruent and incongruent, and calculated separately in the two contexts. Consistent and exploratory trials were defined as when all four moving averages were above and equal or below chance (0.5), respectively. We modelled choices of mice during consistent trial blocks with Bernoulli distributions: p=1 for the contextually correct modality go signal. Opposite models parameterized the distribution with *p*=1 for auditory signals in visual context and vice versa. We employed a baseline random lick model with a bias so that we set *p* to the frequency of lick choices. Then a context-aware model was also constructed, with the *p* equal to the probability of the go instruction, and either a bias, *β*, or a lapse, *λ*, parameter, so that p equals either 1+*β* and *β*, or 1-*λ* and *λ*, for go and nogo trials respectively. The p values of all models were clipped between 0.001 and 0.999. Models were compared during consistent incongruent trials by their mean log likelihoods.

## Electrophysiology

Extracellular multielectrode arrays were manufactured using the same process described previously (Shobe et al., 2015). Each probe had 2 shanks with 64 electrode contacts (area of each contact 0.02 μm2) on each shank. Each shank was 1.05 mm long and 86 μm at its widest point and tapered to a tip. Contacts were distributed in a hexagonal array geometry with 25 μm vertical spacing and 16-20 μm horizontal spacing), spanning all layers of the cortex. Each shank was separated from the other 400 μm. The electrodes were connected to a headstage (Intan Technologies, RHD2000 128-channel Amplifier Board with two RHD2164 amplifier chips) and the headstage was connected to an Intan RHD2000 Evaluation Board, which sampled each signal at a rate of 25 kHz per channel. Signals were then digitally band-pass-filtered offline (100 – 3000 Hz) and a background signal subtraction was performed (Shobe et al., 2015). To ensure synchrony between physiological signals and behavioral epochs, signals relevant to the behavioral task (licking, water delivery, visual/auditory cue characteristics and timing, and locomotion) were recorded in tandem with electrophysiological signals by the same Intan RHD2000 Evaluation Board.

## Microprobe implantation

On the day of the recording, the animal was first handled and then the headbar was attached to head-fix the animal on the spherical treadmill. The elastomer sealant Kwik-Sil was removed and cortex buffer (135 mM NaCl, 5 mM KCl, 5 mM HEPES, 1.8 mM CaCl2 and 1 mM MgCl2) was immediately placed on top of the craniotomy in order to keep the exposed brain moist. The mouse skull was then stereotaxically aligned and the silicon microprobe coated with a fluorescent dye (DiI, Invitrogen), was stereotaxically lowered using a micromanipulator into the ACC to a depth of 0.85 mm. This process was monitored using a surgical microscope (Zeiss STEMI 2000). Once inserted, the probe was allowed to settle among the brain tissue for 1 hr. Recordings of multiple single-unit firing activity were performed during task engagement (approximately 1 hr). After the recording, the animal was anaesthetized, sacrificed, and its brain was extracted for probe confirmation.

## Single unit activities

(SUA). Spike sorting was performed by Kilosort 2 (Pachitariu et al., 2016), and then manually curated in phy2 using MATLAB and PYTHON yielding single unit activities. Standard consistency criteria were employed for autocorrelograms, interspike interval histograms, waveforms, cluster split and merge, maximal amplitude electrode locations, low false positive or missed spikes and also stable feature projections throughout the recording session. We only included units without long-timescale drift, corresponding to unit separability in feature space, stationarity of signal to noise ratio (S/N), and stationarity of firing rate.

## Spike counts

Spike counts were calculated in 10-ms sliding windows, then Gaussian-smoothed (σ=100 ms), approximates single trial instantaneous firing rates (IFR). For decoders IFRs were transformed to z-scores, with mean calculated from prestimulus (from -1500 ms to 0 ms) time-averaged baseline activities, while standard deviation was calculated for the whole trial.

## Geometry of neural representations

We regarded neural activity as a time-varying vector of baseline-standardized IFRs. The components constitute a basis for the population activity vector space, the possible activity patterns of the neuron population. Classification of trials between values of task variables was trained with logistic regression. Test results were shown as 10 fold cross-validation mean over folds. At each time point we trained separate decoders. The decision vector (DV) is the normal vector along which the most difference between classes can be found in trial to trial variation. Angles between decoders were calculated as γ_12_ = arccos d_1_ d_2_, normalized at all timepoints. DVs of multiple decoders with the same underlying neural basis define low dimensional subspaces of task relevant activity. Two-dimensional subspace of two decoders was visualized by the first DV as the first coordinate, and the second DV projected with QR decomposition to the closest orthogonal line to the first DV as second coordinate. Effective chance level was averaged over 40 independent decoder cross-validation accuracy distributions, with fully randomized trial labels from a Bernoulli (p=0.5) distribution. Threshold consisted of s.e.m. from 40 random samples plus from CVs, with one-sided confidence-level.

Animal experiments and analysis are further detailed in (Hajnal et al., 2021).

## Recurrent Neural Network Model

A standard recurrent neural network (RNN) model without gates was built to process the same stimuli and trial structure the mice were tasked to: Visual and auditory stimuli were either ‘go’ or ‘no go’, and a decision was to be made during stimulus presentation. Context determined which modality to base the decision on. The decision was compared to the contextual target, and either rewarded or punished for success or failure respectively. The RNN was governed by the following sequential (recursive) update equations:

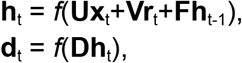

where **x** is the multidimensional combined audio and visual input, **r** is a maintained reward variable for the outcome in the previous trial (not the previous timestep), **h** is the hidden layer activity vector variable, **d** is the output vector variable, with *t* and *t-1* timesteps indices. **F** is the recurrent connections for the hidden layer, while **U, V**, and **D** are the stimulus, reward input map weights, and the output map weights respectively. The non-linearity was chosen *f*=*tanh*, as all variables were one-hot encoded: E.g. **x** = (1, 0, 0, 1) is combined from a (1,0), ‘go’, visual- and (0,1), ‘no go’, auditory-valued vector. The discrete decision was constructed by round(softmax(**d**)), i.e. the component with the larger value was taken as the winner between the two components. The reward was calculated as **r** = (1,0) for success, when the decision equaled the target, and **r** = (0,1) for error trials.

For a single trial, a time resolution of 15 time points was chosen: a 9 point long stimulus presentation with 3 pre and 3 post stimulus points. As opposed to the mouse experiment, we omitted timeout on errors. The network had to decide on the response during the stimulus presentation at time point 7 from stimulus onset, and after the 7th point received reward or punishment until the next trial’s stimulus began. We found this time resolution as optimal: Oscillations were typically not present in the hidden activity, but the time course was detailed enough to observe dynamical changes.

A single data point consisted of 5 subsequent trials, each with 15 timepoints, from the same context in a sequence: 4 trials as a batch of trials during which the model had the opportunity to infer context, and the final trial where the loss function was calculated from the last decision as input to gradients for backpropagation. We generated data combinatorially for 2 choices by 2 modalities in 5-trial sequences, reaching 1024 different sequences total per context. The entire 2048 long dataset of 5-trials sequences was used as training. We did not perform cross-validation for the following reasons: i) each sequence was unique in the dataset, ii) each sequence that contained at least one incongruent trial was successfully completed; in contrast there were only cca. 3% (32+32) of genuinely unsolvable trials: a sequence type consisting of 4 congruent and 1 final incongruent trial, iii) there was no additional stochasticity in the data to reduce the role of deterministic context inference.

We assessed the network performance for the number of hidden neurons from 3 to 100, and found that the smaller the network size, the more random initialization determines the learning capacity of the network within reasonable training iterations. We settled at the minimum feasible hyperparameter set: hidden layer size 30, with 5000 training epochs and used the ADAM optimizer with initial learning rate at 10^−4^. We trained 100 models with different, fixed random generator seeds.

## Analysis of the RNN performance

Linear decoders for the context variable were constructed with 10-fold stratified cross-validation, the accuracy estimate from each fold averaged, and the s.e.m. calculated. Decoders were calculated at each timepoint throughout the 5-trial sequence. Fraction correct estimates for the network outputs were calculated at each decision requirement point for each of the 5 trials in the sequence. Hidden layer neural activity trial-averages were constructed for stimuli variables. Decoders, fraction correct estimates, and trial-averages were also made available to be calculated from designated subsets of trials, such as incongruent trials, or sequences of specific patterns.

## Data availability

Behavioral and spike sorted electrophysiological data, and pretrained RNN models are available in the Zenodo database: https://zenodo.org/record/8379272

## Code Availability

Event generator and behaviour data acquisition: https://github.com/GolshaniCHS/AttentionTask

Analysis: https://github.com/CSNLWigner/mouse-acc-rnn-contextgatedattention

## Notes

https://zenodo.org/record/8379272

### Competing Interest Statement

The authors have declared no competing interest.

